# Whole-genome resequencing revealed the Origin and Domestication of Chinese Domestic Rabbits

**DOI:** 10.1101/2024.03.26.586758

**Authors:** Kerui Xie, Zichen Song, Yanyan Wang, Yan Di, Wenqang Li, Yubin Wang, Aiguo Yang, Xibo Qiao, Bo Wang, Mingyong Li, Xiping Xie, Xiaohong Xie, Lie Liu, Chao Ning, Hui Tang, Xianyao Li, Xinsheng Wu, Qin Zhang, Dan Wang, Xinzhong Fan

## Abstract

The evolutionary and genetic origins of Chinese indigenous rabbits (Oryctolagus cuniculus) remain largely unclear, despite being among the most recently domesticated animals. We sequenced the whole genomes of 142 individual rabbits and collected 25 resequencing accessions from the NCBI, representing six Chinese indigenous breeds, three other domesticated breeds (serving as a bridge between European wild-type and Chinese domestic populations), and two wild rabbit populations from the Iberian Peninsula and Southern France. Population and demographic analyses suggest that Chinese domestic rabbits are most likely descendants of O. c. cuniculus, native to France 800-1500 years ago. These rabbits likely first arrived in the southeast coastal areas of China through trade before spreading to inland regions. Additionally, there may be other origins for Chinese domestic rabbits. We observed considerable variation in the genetic makeup of maternal ancestry between Chinese domestic rabbits and European wild populations, with Chinese rabbits possessing unique mitochondrial haplotypes. Our analysis also highlights selective sweeps on genes affecting brain and neuronal development, which may have been under strong positive selection during domestication; genes related to starch digestion and fat metabolism, suggesting an evolutionary adaptation to digest high-starch diets; and the white coat phenotype in rabbits, resulting from selection at the melanogenesis-associated transcription factor locus. Overall, our data provide comprehensive insights into the origin and domestication of rabbits and lay the foundation for genome-based breeding.

## Introduction

The domestic rabbit (Oryctolagus cuniculus) stands out as a unique and multifaceted domesticated species. It includes many different breeds, strains, and populations, featuring unique phenotypic variations and extremely diverse performance and production characteristics. significantly impacting the economy and playing a pivotal role in its ecological niche alongside wild and feral counterparts [1]. It boasts a broad geographic distribution, serving as valuable livestock for meat and fur production and playing a crucial role as an animal model in biomedical research [2, 3]. It is the sole species that has been domesticated in its family. This Leporidae animal consists of two recognized subspecies, *O. cuniculus algirus* and O. cuniculus cuniculus[4], which diverged approximately 1.8 million years ago [5]. O. c. algirus is found in the southwestern part of the Iberian Peninsula, while O. c. cuniculus is located in the northeastern Iberian Peninsula and France [6].

The rabbit is also called the European rabbit because of its origin in Europe. Its domestication is mainly believed to have occurred around the 5th century, primarily on the Iberian Peninsula and in southern France. Monks in southern France played a significant role in the early domestication of rabbits, breeding them for various purposes, including as a food source during Lent [7]. During the Middle Ages, rabbits were extensively relocated by humans, and today they have become one of the most widely distributed mammals, found across all continents and hundreds of islands [8, 9].

There is a long history of rabbit domestication and breeding in China. Historical records and cultural references indicate that rabbits have been raised in China for various purposes, including as pets, for meat, and for fur for centuries [10]. The small size and fragile bone structure of rabbits lead to the rarity of preserved rabbit remains at archaeological sites, often due to predation or microbial decomposition, so it is difficult to yield information about rabbit remains found in ancient tombs. However, there are artifacts related to rabbits in China [10]. Notably, China has several rabbit resources and is recognized as a major rabbit breeding country, having the largest rabbit population in the world [11]. The significant role of rabbit and its contribution to the Chinese economy underscores the importance of rabbits in Chinese history and culture. However, for detailed information on the origin of domestic rabbits in China, there are different insights. One main insight is the “European Origin” theory. It suggests domestic rabbits were first domesticated in Europe. The results are there is no wild rabbits (belong to the genus Oryctolagus) but wild hares (belong to the genus Lepus) located in China now, and no fossils of rabbits have been discovered in China or anywhere in Asia, suggesting that these burrowing rabbits were introduced from outside the continent [12]. Another insight is the “Local Origin” theory for domestic rabbits proposes that rabbits were domesticated independently in various regions rather than having a single origin. Historical records do mention Himalayan rabbits located in the Himalayan region, and other rabbit breeds are not adapted to the geographic environment in this area [13, 14]. The case supports the indigenous origin of rabbits. Yet, this rabbit breed is also referred to as the Russian, Polish, and Egyptian rabbit, creating ambiguity about its true origin in the international context [15]. Zhao et al., utilizing RAD sequencing technology, investigated the genetic links between Chinese and European rabbits, proposing that China’s domestic rabbits might have an indigenous origin and domestication history, independent of European influences [16]. Contrasting this, Long et al. revealed limited genetic diversity in the mtDNA D-loop of Chinese rabbit breeds, notably poorer than that of European breeds, suggests a European origin for China’s domestic rabbits [17]. Consequently, the origin of China’s domestic rabbits remains a subject of debate.

To comprehensively understand the origins, domestication, and diversification of Chinese indigenous rabbits, we collected resequencing data from 167 representative individuals of domestic rabbit breeds from China, Europe, and the United States, identifying approximately 18.85 million SNPs and 2.12 million insertions-deletions polymorphisms (InDels; <50 bp). Our integrated analysis of mitochondrial (MT) and nuclear genome data supports European origin for the Chinese indigenous rabbit, which was independently domesticated from Oryctolagus cuniculus, and evolved new breeds through genetic mutations during its spread in China. The earliest formation of indigenous rabbits in China dates back 300-800 years ago in the Fujian region, from where they spread to other parts of China through trade. Moreover, we conducted selective scans to elucidate the characteristics formed during the domestication process include changes in behavior patterns and coat color. We detected specific selection in genes related to the nervous system and brain development in domestic rabbits, which may help explain the genetic basis of their adaptation to human environments, enabling them to thrive in enclosed spaces and under human care. We also identified genes associated with fur color, vision, and digestion, reflecting convergent patterns in the domestication process. Additionally, various genes related to body size and reproduction were identified, facilitating integrated practices in selective breeding combined with genome-wide association studies (GWAS).

## RESULTS

### Genetic variation and population structure

We resequenced 142 rabbits (Fig. 1A), including 92 Chinese indigenous rabbits (Sichuan White [SCW] n = 20, Laiwu Black [LWB] n = 20, Jiuyishan [JYS] n = 14, Fujian White [FJW] n = 18, Minxinan Black [MXN]), 50 imported breeds (introduced rabbits Angora [AN] n = 20, Rex [REX] n = 15, and Kangda Meat [KD] n = 15). Additionally, we obtained data from 25 rabbits, including 15 wild rabbits (*O. c. cuniculus* and *O. c. algirus* [WILD] n = 15), Three of them come from the south of Southern France, and the rest come from Iberian Peninsula. and 10 introduced rabbits (New Zealand White [NZW] n = 10), from the NCBI database (Supplemental Table S1). A total of 1168.47 Mb of paired-end reads were generated, with a read length of 150 bp per read. Rabbit genome alignment showed an average depth of 15.31X per individual. Following quality control on the raw data, each individual obtained an average of 146 million high-quality reads, which corresponded to approximately 21.84 Gb of sequenced bases. The average Q20 and Q30 scores were 97.10% and 91.61%, respectively, indicating high sequencing quality that met subsequent analysis requirements (Supplemental Table S2). A total of 18,848,966 high-quality population single nucleotide polymorphisms (SNPs) and 4,590,856 insertions/deletions (INDELs) were identified across the samples (Fig. 1B and Supplemental Table S3). 111,976 SNPs (0.60%) and 5,462 InDels (0.11%) located in the coding regions. Among them, 27,966 SNPs (1.48%) cause codon changes, transcript elongation, or premature termination codons, 2,554 InDels (0.06%) lead to frameshift mutations (Supplemental Table S4 and 5).

**Fig. 1.**
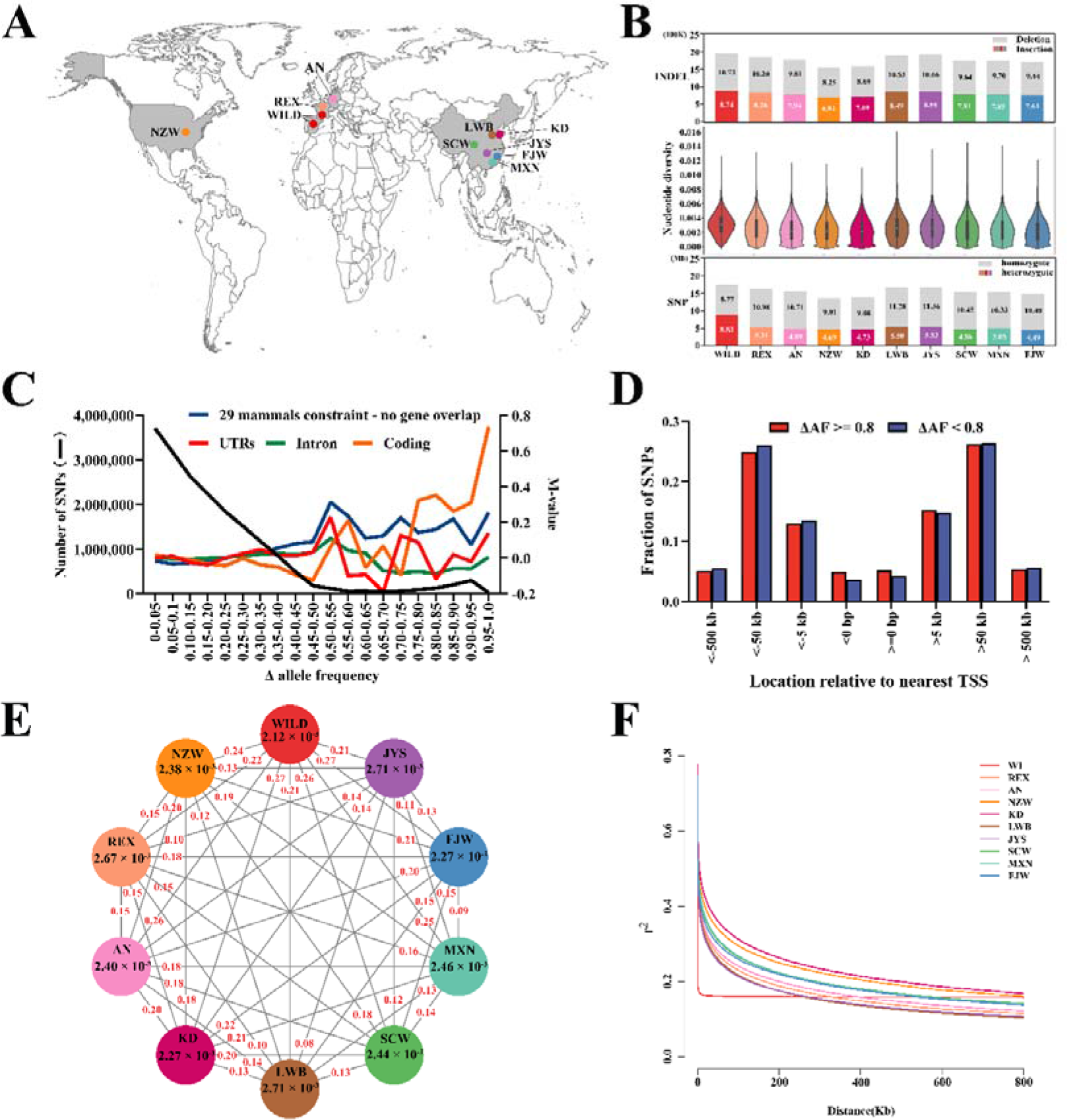
Experimental design and variants statistics. (A) Geographic distributions of 167 rabbit accessions (Sichuan White [SCW]; Laiwu Black [LWB]; Jiuyishan [JYS]; Fujian White [FJW]; Minxinan Black [MXN]; Angora [AN], the origin is in France, and now it is being bred in Shandong; Rex [REX], the origin is in German, and now it is being bred in Shandong; Kangda Meat [KD]; O. c. cuniculus and O. c. algirus [WILD]). The geographic map was drawn using R ggplot2. (B) Genomic variation of 10 populations. Mean number of SNPs and heterozygous and homozygous SNP ratio in the 10 populations are shown at the bottom. Nucleotide diversity ratios of the 10 populations are shown at the middle. Number of insertions and deletions in the 10 populations are shown at the top. (C) The majority of SNPs showed low DAF between wild and domestic rabbits. The black line indicates the number of SNPs in nonoverlapping DAF bins (left y axis). Colored lines denote M values (log2-fold changes) of the relative frequencies of SNPs at noncoding evolutionary conserved sites (blue), in UTRs (red), exons (yellow), and introns (green), according to DAF bins (right y axis). M values were calculated by comparing the frequency of SNPs in a given annotation category in a specific bin with the corresponding frequency across all bins. (D) Location of SNPs at conserved noncoding sites with DAF ≥ 0.8 SNPs and DAF< 0.8 SNPs in relation to the TSS of the most closely linked gene. (E) F coefficient estimates (i.e. (<observed hom. count> - <EXPECTED count>) / (<TOTAL observations> - <EXPECTED count>) and population divergence (FST) across the ten groups. The value in each circle represents a measure of F coefficient estimates for each group; values in red on each line indicate pairwise population divergence between groups. (F) LD decay plots of ten duck populations.

Highly differentiated single SNPs are likely to be direct targets of selection or occur near sites under selection [18]. We identified that 60.8% of the SNPs are located in intergenic regions, while only 0.6% are located in exonic regions. We identified a total of 41,147 non-redundant coding SNPs from ten populations, 66.5% of the SNPs are synonymous SNV (Supplemental Table S5). We calculated the absolute allele frequency difference (ΔAF) between wild rabbits and domestic rabbits, and divided it into 5% bins (ΔAF = 0 to 0.05, etc.). Most SNPs exhibit low ΔAF between wild rabbits and domestic rabbits (Fig. 1C). We examined exonic, intronic, UTR, intergenic, and evolutionarily conserved sites to identify enrichment of high-ΔAF SNPs, as would be expected under directional selection with many independent mutations (Supplemental Table S6). In the intronic, we observed no consistent enrichment of high ΔAF SNPs. However, we found significant enrichment in exonic and conserved non-coding sites (χ2 test, P < 0.05). We found that in each partition with ΔAF > 0.25, there was a significant excess of SNPs in conserved non-coding sites (χ2 test, P < 0.05). However, in coding sequences, a significant excess was only observed when ΔAF > 0.75. Compared to the relative proportions in the entire dataset, there were 4,724 additional SNPs in conserved non-coding sites with ΔAF > 0.25, while only 1351 additional SNPs were observed in exonic regions with ΔAF > 0.75 (Table S6). Therefore, in the domestication process of rabbits, changes in regulatory sites have played a more significant role in terms of quantity than changes in coding sequences. We selected 12,610 SNPs with ΔAF > 0.80 in conserved non-coding regions, representing 1,538 non-overlapping 1-Mb blocks in the rabbit genome. To avoid bias caused by strong linkage disequilibrium, we chose only one SNP every 50 kb [18], resulting in a final set of 7,100 SNPs. Over 60% of these SNPs were located at least 50 kb away from the nearest transcription start site (TSS) (Fig. 1D), indicating that many divergent SNPs are found in long-range regulatory elements.

In addition, we also calculated the nucleotide diversity, wild rabbits had higher nucleotide diversity (πC=C3.1C×C10−3) compared to domesticated populations (πC=C2.4C×C10−3) (Fig. 1B). Generally, domesticated populations have fewer SNPs (*t* test, *P* < 4.42 × 10^-4^) and lower nucleotide diversity (*t* test, *P* < 3.05 × 10^-6^, Supplemental Table S7). We calculated the degree of heterozygosity. Domestic rabbits have a lower heterozygosity of 0.26 compared to wild rabbits with a heterozygosity of 0.34 (Supplemental Table S7), and strong genetic divergence from the other 9 groups (pairwise F_ST_C≥C0.21; Fig. 1E). Moreover, linkage disequilibrium (LD) decayed faster in wild rabbits than in domesticated rabbits (Fig. 1F), indicating a lower degree of genetic recombination in the wild rabbits. This could be associated with the higher levels of artificial selection and inbreeding within the domestic rabbit population.

Next, we analyzed the genetic structure of the rabbit population for clusters (K) ranging from 2 to 6, based on 18,848,966 SNPs among the 167 rabbits. With K = 2, a clear division was found between wild-type rabbits (WILD) and domesticated rabbits (LWB, SCW, JYS, FJW, and MXN). The domestic rabbits shared corresponding ancestral components to the wild rabbits in the French region, which is consistent with the findings indicating that the domestic rabbit originated in France [19]. With K = 3, a clear division was found between Chinese indigenous breeds (FJW, MXN, JYS, LWB and SCW) and European lineage (REX, AN, NZW, and KD). When K = 6, clusters maximized the marginal likelihood (Supplemental Fig 1). The domestic populations shared the main ancestral compositions with their wild counterparts occupied in French (*Oryctolagus cuniculus cuniculus*), with one genetic makeup absent in the domestic population. Likewise, the Chinese indigenous breeds were different to the Europe populations, but they were more similar compared to the French wild population. In addition, the bloodline composition of REX, LWB, and JYS is complex, which may be due to extensive gene flow between them and other rabbit populations. (Fig. 2A). Both phylogenetic relationships, based on a neighbor-joining tree of pairwise genetic distances of whole-genome SNPs and principal component analysis (PCA), indicated three distinct clades. Clade І consisted only of wild-type rabbits. Clade II included introduced breeds and Chinese indigenous breeds (Fig. 2B).

**Fig. 2.**
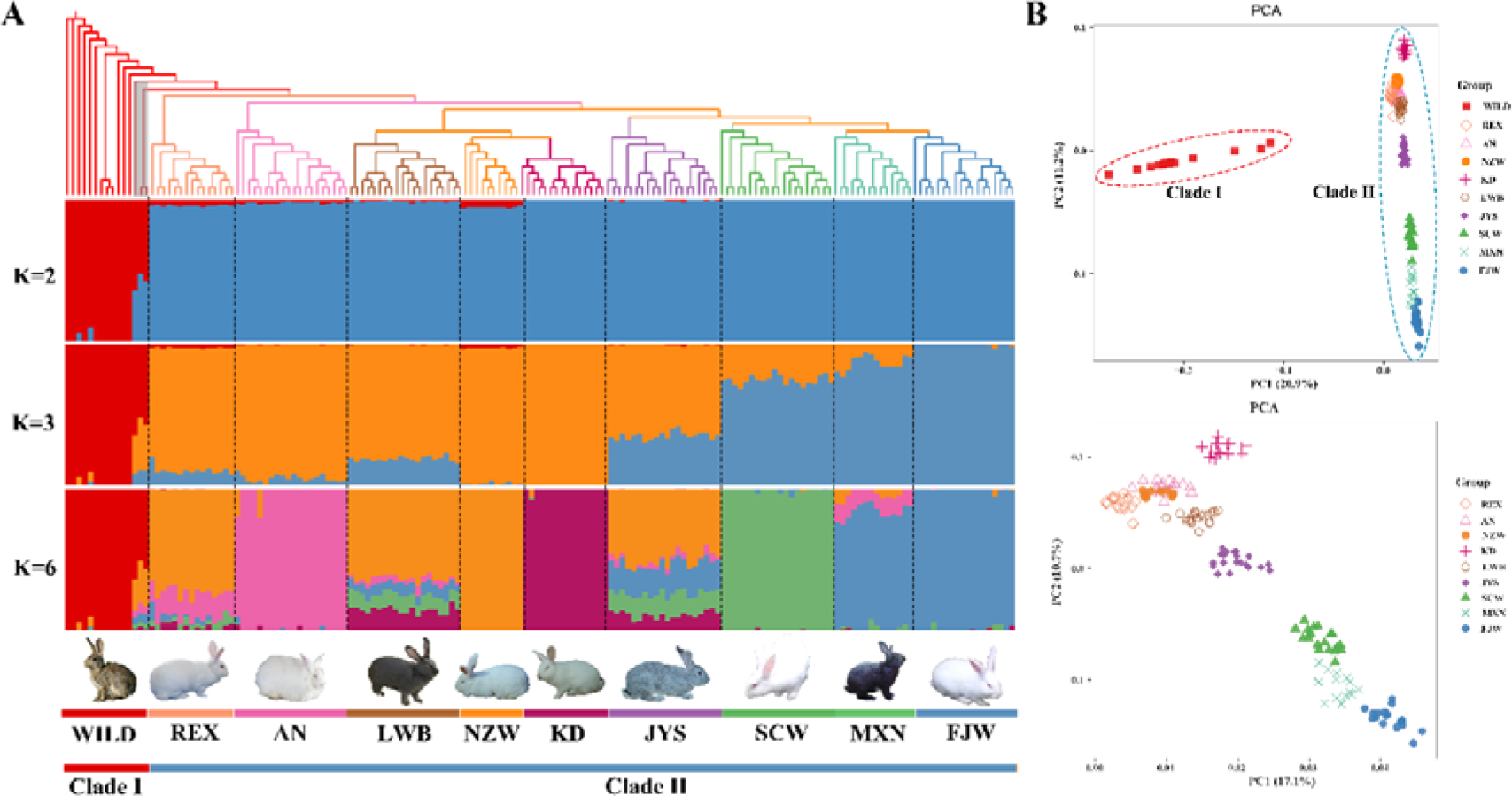
Population structure. (A) The maximum-likelihood phylogeny of 167 rabbit accessions at the top. Population genetic structure of 167 rabbit. At the middle, the length of each colored segment represents the proportion of the individual genome inferred from ancestral populations (K = 2, 3, 6). The population names and clade type are at the bottom. (B) PCA plot of all rabbit populations at the top. Eigenvector 1 and 2 explained 20.9% and 11.2% of the observed variance, respectively. PCA plot of domesticated rabbit populations at the bottom.

### Domestication and dissemination of Oryctolagus cuniculus

To estimate the trajectories of changes in the effective population size (Ne) of ancestral populations of Chinese indigenous rabbits, we conducted a pairwise sequentially Markovian coalescent (PSMC) analysis among Chinese indigenous rabbits and transitional breeds. The populations showed consistent trends with two expansions and two contractions in their effective population size (Ne). One notable expansion was observed during the Penultimate Glaciation Period (0.30-0.13 million years ago) and the Last Glacial Period (110–12 thousand years ago [20]) in each population, followed by subsequent contractions (Fig. 3A). To investigate near-term changes in effective population size over time in each population, we used SMC++ for population history inference. Around 1000 years BP, the Ne model of domestic rabbits was relatively consistent. since 500 years, the domestic breeds have gradually expanded, with FJW of a Chinese indigenous breed and AN, REX, and NZW of transitional breeds increasing consecutively, and other domestic breeds expanding latterly within the last 200 years (Fig. 3B).

**Fig. 3.**
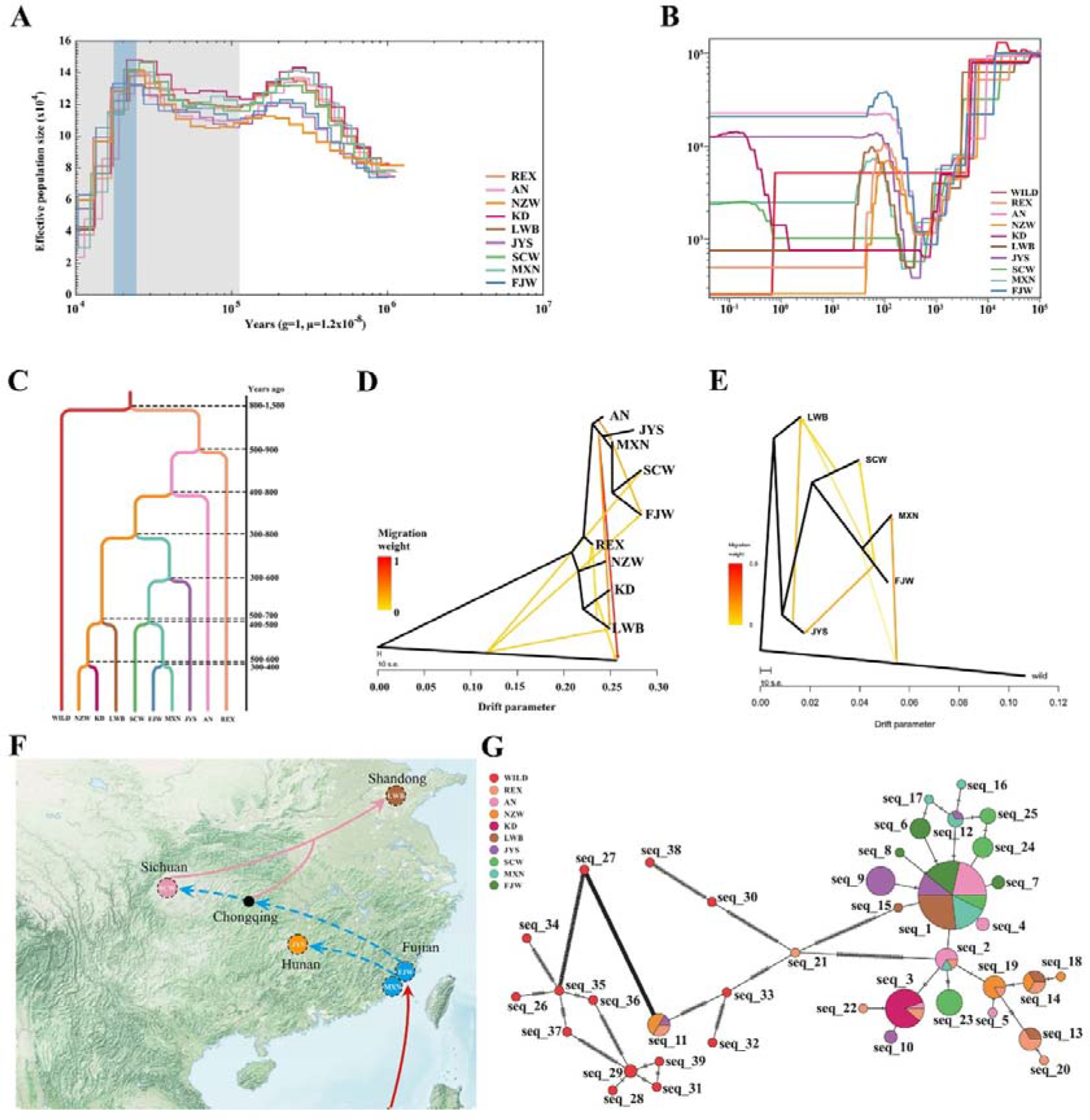
Speciation and demographic history of rabbit. (A) Examples of PSMC estimate changes in the effective population size over time, representing variation in inferred Ne dynamics. The lines represent inferred population sizes and the gray shaded areas indicate the Pleistocene period, with Last Glacial Period (LGP) shown in darker gray, and Last Glacial Maximum (LGM) shown in light blue areas. (B) Examples of SMC++ estimate changes in the effective population size over time. (C) Divergence time for ten groups was estimated using SMC++. (D) Detection of gene flows among all rabbit groups by TreeMix analysis. Arrows represent the direction of migrations. Horizontal branch length is proportional to the amount of genetic drift that has occurred on the branch. Scale bar shows ten times the average standard error of the entries in the sample covariance matrix. (E) Detection of gene flows among Chinese indigenous rabbit by TreeMix analysis. (F) Putative propagation routes of Chinese indigenous rabbit. The dashed lines represent inferences made by combining results from papers, while the solid lines are recorded in the literature. ①The Fujian white rabbit arrived in China via shipping [24]. ② Records of white rabbits being introduced to Chongqing suggest, based on the estimated time of divergence, that they may have come from Fujian [26]. ③The Central Plains region began to introduce white rabbits during the Chongzhen period, and some of these white rabbits came from Sichuan [26]. The geographic map was adapted from NASA (https://visibleearth.nasa.gov/images/147190/explorer-base-map/147191w/). (G) Detection of gene flows among Chinese indigenous rabbit groups by TreeMix analysis.

Next, we employed the diffusion approximation method for the allele frequency spectrum (∂a∂i) to test various demographic scenarios related to domestication. Among the 34 divergent models tested (https://github.com/dportik/dadi_pipeline/), the model of divergence of domesticated breeds (Adjacent secondary contact, longest isolation) was consistent with our population structure results and had the lowest Akaike Information Criterion (AIC) value, indicating a better overall fit with the data (log-likelihood = -10,099,671.73; AIC = 20,199,363.46) (Supplemental Fig 2, Supplemental Table S8). France *Oryctolagus cuniculus* differentiated from the Iberia population [18] (Supplemental Fig 3), was subsequently domesticated into the domestic rabbit, and then domestic rabbits formed the current introduced breeds and local Chinese indigenous breeds through human trade and selection. In addition, we also found gene flow between the wild rabbit population and the domestic rabbit population, which is consistent with the laws of domestication of domestic animals. To delineate domestication and dissemination, we further constructed the phylogenetic relationships between them and used SMC++ to calculate the specific divergence times for each population (Fig. 3C). The combination of the PSMC results supports the hypothesis of a common ancestor between Chinese indigenous breeds of rabbits and introduced breeds. Approximately 800-1,500 years ago, domestic rabbits diverged from their wild counterpart. We speculate that *Oryctolagus cuniculus* was firstly domesticated in France, based on its current geographic distribution and historical records [19]. The divergence of the ancestors of an imported breed (European domestic rabbit) occurred 400-900 years ago, during the 15th and 16th century, the domestication of rabbits in Europe had reached a mature stage [21]. The divergence time between Chinese domestic rabbits and European domestic rabbits is approximately 300-800 years ago. During this period, there is close trade interaction between China and Europe [22]. In the early 16th century, the Ming Dynasty relaxed its maritime restrictions, allowing Portuguese people to be the first to come to the coastal areas of China. Soon after, Spanish and Dutch people followed suit [22]. Interestingly, it was during the mid-Ming Dynasty in the 16th century that the domestic rabbits began to appear in Chinese historical records. By the late Ming period, there were official records of rabbit breeding in China [23]. In the mid-17th century, there are records documenting the large-scale introduction of white domestic rabbits to Fujian in southeastern China [24, 25]. The white domestic rabbits then spread westwards to Hunan, Guizhou, Chongqing, and later entered Sichuan before finally reaching the Central Plain region (now Shandong and Henan provinces) [26] (Fig. 3F).

Multiple gene flows were detected among the ten groups by Treemix analysis (Fig. 3D). The gene flow from KD and AN to WILD can be attributed to domesticated rabbits escaping into the wild during the taming process. Additionally, genetic exchange occurred between wild rabbits and native rabbit populations in China. This was likely facilitated by the transportation of wild rabbits to China on multiple occasions. The proportion of wild rabbit introgression fragments detected in domestic rabbits varied from 0.88% to 0.38%, averaging 0.6%, We also found that the proportion of wild rabbit introgressed fragments in France is significantly higher than that in Iberia (Supplemental Fig 4), which aligns with the domestication of rabbits in France. There is also significant genetic exchange between introduced breeds and indigenous Chinese rabbits, likely due to human efforts to improve the production performance of domestic rabbits through crossbreeding with other rabbit breeds. The most significant characteristic of Chinese indigenous rabbits, as opposed to introduced breeds, is their small size, slow growth, strong resistance, and excellent meat quality. This may base attributed to differences in the domestication process when compared to European rabbits. However, the growth rate and body size of the LWB surpass that of other indigenous domestic rabbits. Considering the gene flow and structural analysis (Fig. 2A), LWB has a closer relationship with meat rabbits. Gene flow Among Chinese domestic rabbit populations showed that FJW ancestors hybridized most frequently with other indigenous breeds, which further supported the speculation that Chinese domestic rabbits originated and spread from Fujian (Fig. 3E).

We investigated the maternal origin of Chinese domestic rabbits by sequencing the mitochondrial genome of the 10 rabbit populations. Totally a total of 39 mitochondrial haplotypes were identified (Supplemental Table S9 and S10). Wild rabbits have highest number of haplotypes representing the most genetic diversity. Wild rabbit mitochondrial haplotypes are absent in domesticated populations, only two domesticated haplotypes show a relatively close genetic affinity to wild rabbits (seq_11 and seq_21, these haplotypes primarily belong to European lineage breeds), and most wild rabbit haplotypes exhibit a high degree of variation compared to domesticated rabbit (Fig. 3G). The main haplotypes are sep_1 (including MXN, FJW, SCW, LWB, JYS and AN). Most sequences were closely related to seq1, differing only by a few substitutions. Chinese indigenous rabbit shares some haplotypes with introduced breeds (seq_2, seq_11, seq_13 and seq_14), indicating a common maternal origin. However, Chinese indigenous rabbits had unique haplotypes, which may have arisen from mutations during breed formation or from a maternal origin distinct from that of other domestic rabbits.

### Adaptation to living environment during rabbit domestication

In order to detect the signature of selection for environmental adaptability during rabbit domestication, we conducted a genome-wide scan for regions exhibiting high nucleotide differentiation (θπ) coefficients across populations of wild and domesticated rabbits. This analysis was performed using sliding windows of 50 kb, and global F_ST_ values were calculated for each population (Fig. 4A). Due to the complicated and not fully resolved demographic history of these populations, it is challenging to establish a clear threshold that can differentiate true selective sweeps from regions of homozygosity caused by genetic drift. Therefore, we also used pairwise diversity ratio (θπ(wild/domesticated)) to identify genomic regions under putative positive selection during domestication. Our analysis identified 1490 genes that achieved high F_ST_ and θπ scores within the top 5% (Supplementary Table S11), indicating that they underwent strong positive selection during rabbit domestication.

**Fig 4.**
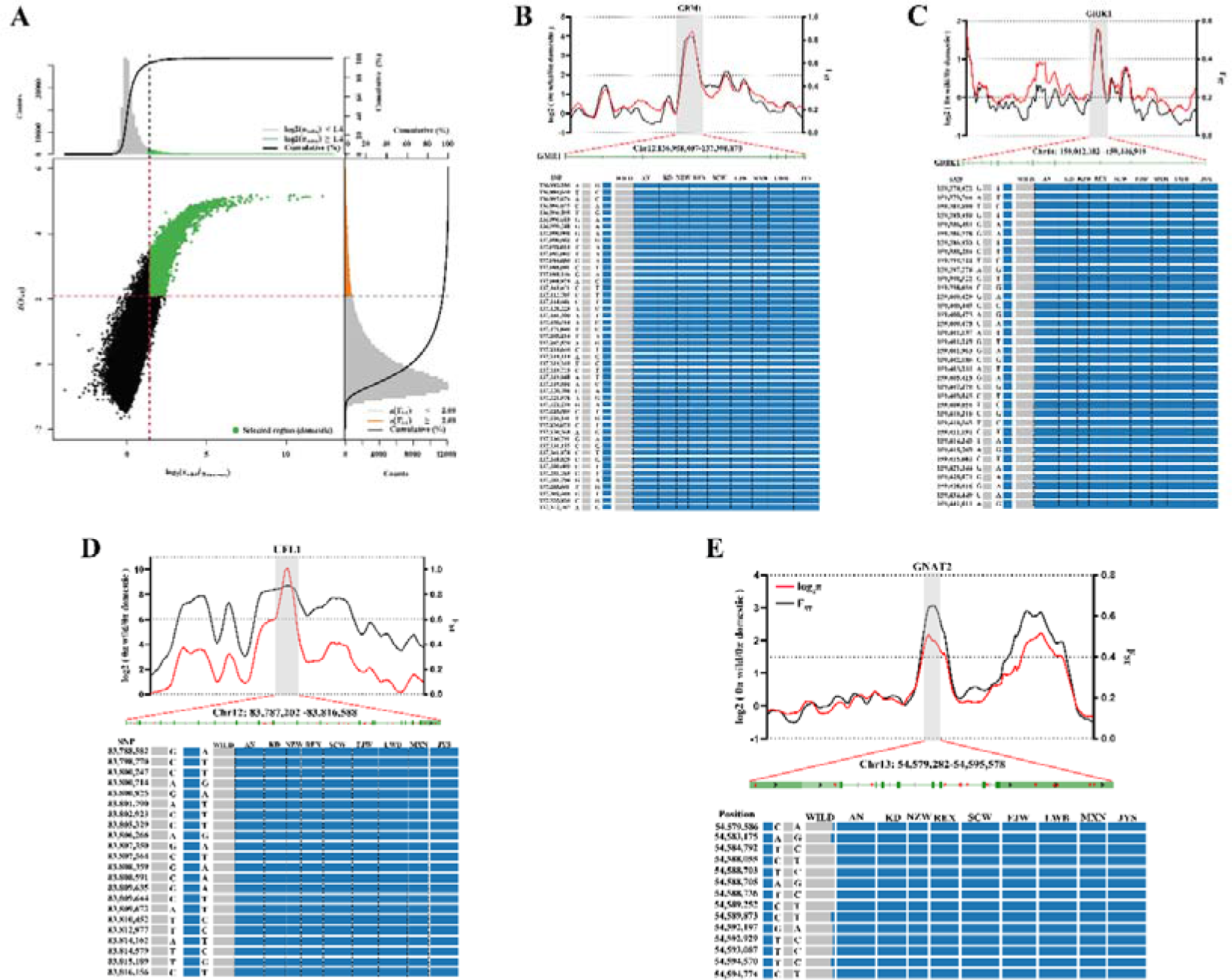
Genomic regions with strong selective sweep signals in wild population rabbits and domesticated population rabbits. (A) Distribution of θπ ratios θπ(wild/domesticated) and Z(F_ST_) values, which are calculated by 50-kbwindows with 10-kbsteps. Green data points located to the top-right regions correspond to the 5% right tails of empirical log_2_ (θπ wild/θπ domestic) ratio distribution, and the top 5% empirical Z(F_ST_) distribution are genomic regions under selection during rabbit domestication. The two horizontal and vertical gray lines represent the top 5% value of Z(F_ST_) (2.09 and log_2_ (θπ wild/θπ domestic) (1.4), respectively. (B) The log_2_ (θπ) ratios and F_ST_ values around the GRM1 locus and allele frequencies of 48 SNPs within the GRM1 gene across ten rabbit populations. The black and red lines represent log2 (θπ wild/θπ domestic) ratios and F_ST_ values, respectively. The gray bar shows the region under strong selection in GRM1 gene. The SNPs were named according to their position on the chromosome. (C) The log2 (θπ) ratios and FST values around the GRIK1 locus and allele frequencies of 36 SNPs within the GRIK1 gene across ten rabbit populations. (D) The log2 (θπ) ratios and F_ST_ values around the UFL1 locus and allele frequencies of 22 SNPs within the UFL1 gene across ten rabbit populations. (E) The log_2_ (θπ) ratios and F_ST_ values around the GNAT2 locus and allele frequencies of 14 SNPs within the GNAT2 gene across ten rabbit populations.

We selected the entire set of 1490 genes located within the top 5% regions for gene ontology (GO) and KEGG pathway analysis. As a result, we obtained a total of 137 enriched GO terms and 24 KEGG pathways (Supplementary Table S12). We focused on GO annotations related to locomotor behavior, neuronal synaptic plasticity, lipid metabolism and energy metabolism, reproduction, and visual perception for the 1490 putative positive selection genes. Among them, we found that entries related to neurons and metabolism are proportionally higher (Supplementary Table S13).

From the highlighted GO terms, 29 neuro-synapse genes were identified as being under positive selection (Supplementary Table S13), with 13 (PLCB1, GRIA4, GNAI3, GLS, GRIK1, GRM1, RAB11A, EPHA7, ADAM22, EPHB1, MAGI2, DISC1, and ALS2) in the top 1% of F_ST_ and θπ (Supplemental Table S14). In particular, GRIK1 (glutamate receptor, ionotropic kainate 1) and GRM1 (glutamate ionotropic receptor kainate type subunit 1) both showed high FST and θπ values compared to neighboring regions, suggesting functional importance. In the GRM1 and GRIK1 gene regions, we identified 48 and 36 loci, respectively, as fixed loci that are homozygous in domesticated breeds and absent in wild populations. This indicates that these specific genetic markers have been selectively fixed through the domestication process, highlighting the genetic differences between domesticated breeds and their wild counterparts (Fig. 4B and C). In addition to neuron-synapse genes, 19 genes have been identified in the pathway related to neural-brain development (Supplementary Table S13), showing high F_ST_ and θπ values. Among these genes, 5 genes rank in the top 1% for both parameters (Supplementary Table S14), including UFL1, in this region, we identified a total of 14 SNPs that are homozygous in domesticated breeds, and absent in wild populations (Fig. 4D).

Beyond the neuronal-synapse-brain genes, 22 genes were identified in the three visual pathways with high F_ST_ and θπ values (Supplementary Table S12). Among these genes, 6 were found with both parameters yielding top 1% ranked values (Supplemental Table S14) such as GNAT2. with 7 of them located in the coding regions. Among these loci, 7 wild rabbits showed partial homozygosity, suggesting that they may be individuals with poorer vision within the wild rabbit population (Fig. 4E).

### Adaptation to a starch-rich diet during domestication of rabbits

Compared to wild animals, domesticated animals have abundant food sources in addition to a safe living environment. In the wild, the main food source for rabbits is plant leaves and stems, which primarily consist of crude fiber. However, under artificial feeding, domesticated rabbits can more easily access food that is high in energy, protein, and carbohydrates. Two amylum digestion and absorption pathways, namely starch and sucrose metabolism and salivary secretion, were found in the selective-sweep signals of domestic rabbits. Four key genes encoding starch-digesting enzymes were identified, namely LOC100342719, LOC100343225, LOC100343481 and LOC100343733. Compared to neighboring regions, they show higher values of F_ST_ and θπ (Fig. 5), indicating their functional significance. they encoded a pancreatic α-amylase to catalyse the first step of starch digestion. Therefore, we decided to investigate the SNP diversity across the entire region to facilitate the identification of causal variations. We found that wild rabbits exhibited much higher heterozygosity in this region compared to domestic rabbits, particularly at the LOC100343733 locus.

**Fig 5.**
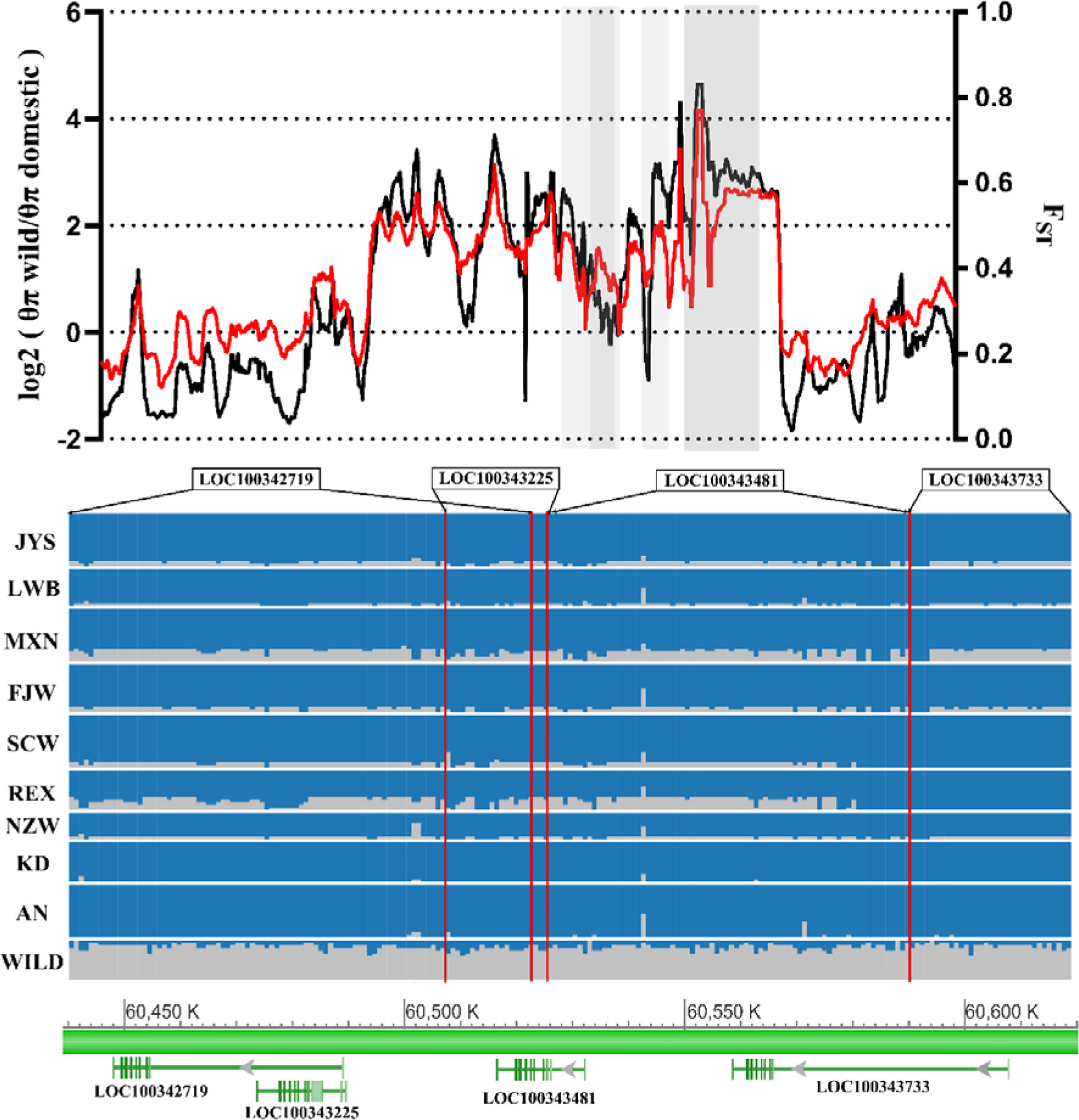
Starch-digesting enzymes gene shows different genetic signatures between wild and domesticated rabbit. The log2 (θπ) ratios and F_ST_ values around the starch-digesting enzymes gene locus and allele frequencies of 205 SNPs within the LOC100342719, LOC100343225, LOC100343481 and LOC100343733 genes across ten rabbit populations. The black and red lines represent log_2_ (θπ wild/θπ domestic) ratios and F_ST_ values, respectively. The gray bar shows the region under strong selection in Starch-digesting enzymes gene. Blue indicates homozygous sites, while gray represents heterozygous sites. The bottom shows the positions of four genes on the chromosome.

### Selection for coat colors

Although the domestication process has resulted in a variety of new phenotypic and behavioral traits, coat color variation is one of the few characteristics that distinguish all domesticated animals from their wild ancestors. In the early stages of domestication, novel coat colors appeared easily and could be selected with little cost to other aspects of the organism [27], often favoring the same variations that are disfavored in nature. In this way, humans satisfied their desire for novelty by selecting for coat colors, sacrificing the survival ability of livestock in the wild. This manipulation of livestock breeding by humans has produced and maintained coat colors that would not be possible in a wild environment [28]. To identify the genetic signatures associated with white coat color, we conducted a genome-wide search for regions exhibiting high FST values between populations of rabbits with white coat (NZW, REX, AN, KD, SCW, and FJW) and those without white coat (WILD, MXN, LWB, and JYS). We have discovered a pathway linked to melanin synthesis in both white-coat and non-white-coat rabbits (Fig. 6A and Supplemental Table S15). Within this pathway, we have identified a highly differentiated region that exhibits overlap with the transcription factor *TYR* (F_ST_ = 0.5) (Fig. 6B). Importantly, this gene holds a pivotal role in regulating melanin production. Within the *TYR* gene region, we found 41 homozygous single nucleotide polymorphisms (SNPs) in all white-coat breeds, whereas all non-white-coat breeds exhibited heterozygosity at these specific loci (Fig. 6C). These mutations were completely associated with the white-coat phenotype, suggesting a causative mutation at the *TYR* locus.

**Fig 6.**
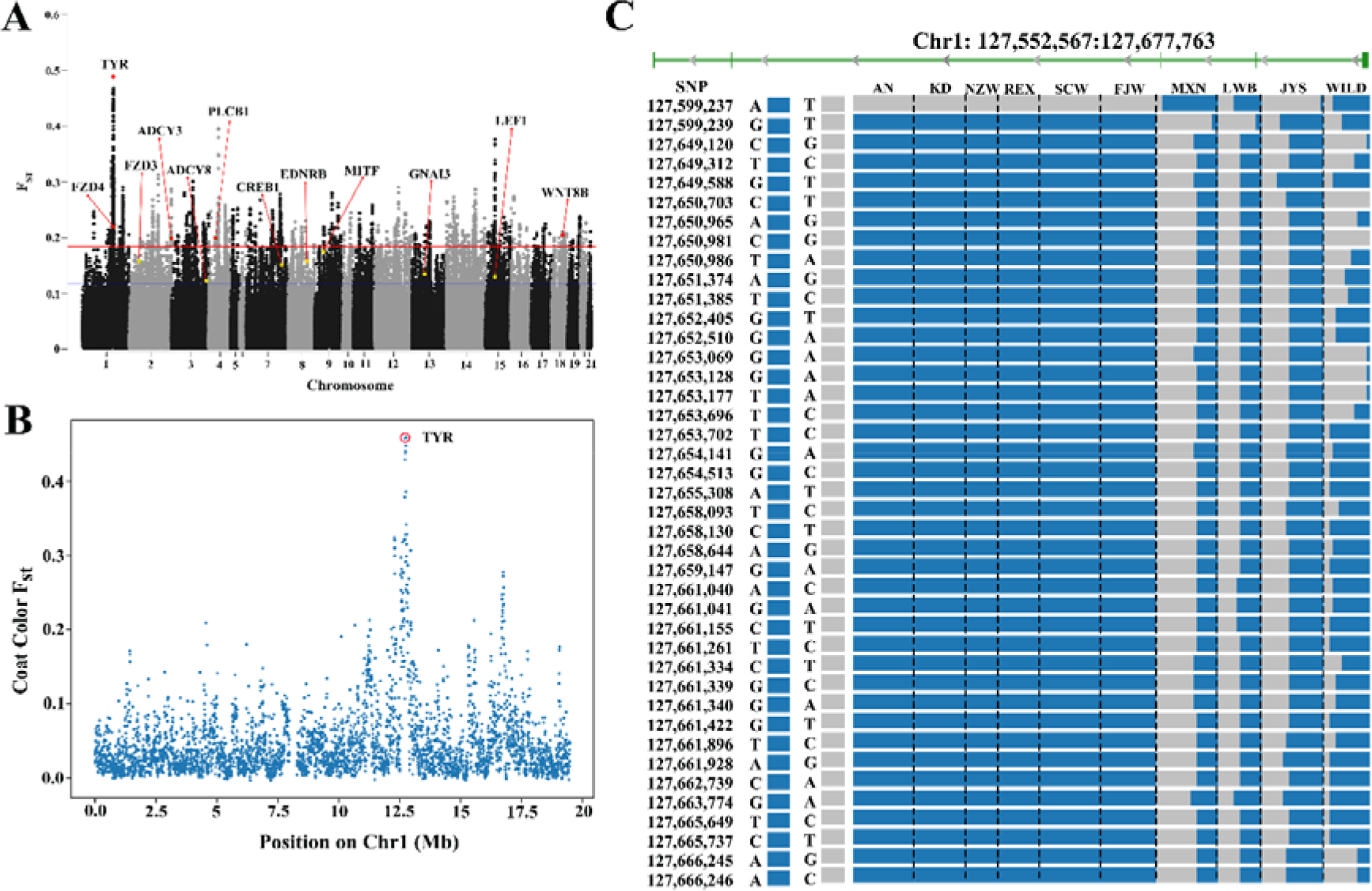
*TYR* shows different genetic signatures between white-coat and non-white-coat rabbit. (A) Genome-wide distribution of selective sweeps related to coat colors in rabbits. (B) F_ST_ plot around the TYR locus. The F_ST_ value of TYR is highest for chromosome 1. (C) The 41 homozygous SNPs in white-coat rabbits and absent in non-white-coat rabbits. SNPs were named according to their position on the chromosome.

**Table 1.**
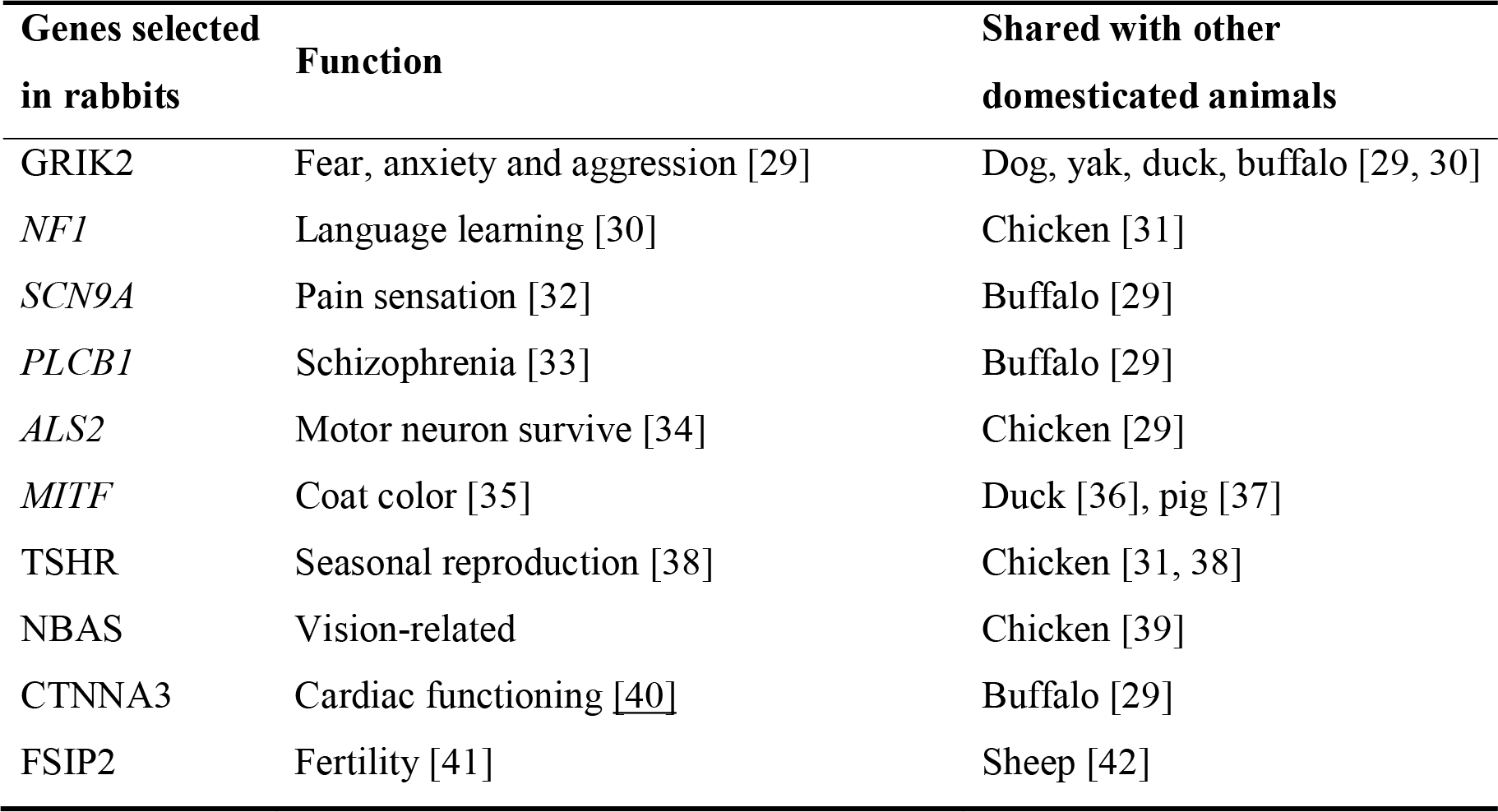
Convergent Genes shared among rabbits and other domesticated animals.

## Discussion

In order to elucidate the origin and domestication of Chinese rabbits, we collected and analyzed the genome data of five Chinese indigenous breeds, four European domestic breeds and two subspecies of wild populations. This pioneering study delves into the genetic architecture, phylogenetic relationships, and origin and domestication process of Chinese indigenous rabbits.

The genomic analysis of Chinese indigenous and European domestic rabbit breeds, alongside two subspecies of wild populations, underscores the complex domestication process of rabbits in China. Our findings suggest that Chinese domestic rabbits likely originated from Oryctolagus cuniculus cuniculus, corroborating previous hypotheses about the domestication route from Europe to China [23]. Notably, the absence of one ancestral composition in domesticated rabbits, which is present in wild French counterparts, points to selective breeding practices that favored certain traits over others. This selective process may have led to the loss of genetic traits found in the wild population, indicating the significant impact of human intervention on the genetic diversity of domestic rabbits [43, 44]. The significant allele frequency differences (ΔAF) between wild and domestic rabbits, especially in conserved non-coding regions, underscore the impact of artificial selection in shaping the rabbit genome. The enrichment of ΔAF in these regions suggests that alterations in gene regulation, possibly affecting traits such as growth rate, size, and fur quality, have been crucial in the domestication process. These findings not only contribute to our understanding of rabbit domestication but also offer a genetic window into the selective pressures and breeding strategies employed by humans over centuries.

Comparison with existing literature reveals that the demographic patterns and divergence times observed in our study align with those of other domesticated animals, suggesting a common response to environmental changes and human activities across species [30, 45–47]. The historical context provided by the decree of Pope Gregory in the 6th century and the subsequent spread of rabbits during the Age of Discovery further enrich our understanding of the domestication timeline [7]. However, the unique haplotypes identified in Chinese indigenous rabbits that cannot be accounted for by European origins hint at a more complex domestication story, possibly involving multiple maternal lineages. This finding challenges the simplistic view of rabbit domestication and opens new avenues for research into the genetic origins and domestication pathways of Chinese indigenous rabbits. Future studies should aim to explore these unique haplotypes further, investigating additional maternal origins and assessing their impact on the genetic diversity and adaptation of Chinese indigenous rabbits. Additionally, a broader genomic analysis incorporating more diverse rabbit populations could shed light on the global patterns of rabbit domestication and genetic exchange.

To identify the genomic regions that underwent selection during domestication, we examined the diversity between wild and domestic samples. Long-term confinement has altered the adaptation of domestic rabbits to the rearing environment. The genes involved in brain and neural development play a targeted role in the domestication process of rabbits. Domestication reduced amygdala volume and enlarged medial prefrontal cortex volume [48]. Three loci, GRIK1, GMR1, and UFL1, exhibited very strong evidence of selective sweeps that are likely associated with domestication. GRIK1 encodes a subunit of a glutamate receptor that plays a role in synaptic plasticity and is crucial for learning and memory processes [49]. GMR1 is responsible for encoding a receptor involved in glutamate metabolism, which plays a role in regulating pain perception and is closely linked to disorders associated with abnormal synaptic plasticity [50]. Additionally, the gene has been associated with depression [51], potentially indicating a connection to the prolonged confinement environment experienced by domestic rabbits. UFM1 is required for animal development and the normal function of multiple tissues and organs. Clinical investigations have detected rare genetic variants in human UFM1 genes that are associated with early-onset encephalopathy and impaired brain development, thus indicating the crucial involvement of the UFM1 system in the nervous system [52].

Vision plays a pivotal role in the natural habitat of animals, influencing crucial survival behaviors such as mating, foraging, and predator evasion [53]. Domesticated animals, protected and provided for by humans, have experienced a reduction in the importance of vision compared to their wild ancestors. Various domesticated species, including dogs, horses, ducks, sheep and chickens, have exhibited varying degrees of diminished visual capabilities relative to their wild counterparts [39, 54–57]. Despite being one of the domesticated species with a recent history of domestication, research on the evolutionary changes in vision during the domestication process in rabbits has been limited. GNAT2 is a critical gene for color vision perception. Wild rabbit populations exhibit heterozygosity, while domestic rabbit populations exhibit homozygosity. Mutations in GNAT2 have been linked to changes in refractive development and increased susceptibility to myopia in mice [58]. Therefore, it is hypothesized that during the early stages of rabbit domestication, there existed both individuals with good vision and individuals with poor vision within wild rabbit populations. The visually impaired wild rabbits were more susceptible to capture and domestication by humans, as well as being easier to manage in later stages of breeding and care. Consequently, these individuals with impaired vision continued to reproduce, resulting in a progressive decline in the overall visual capabilities of domestic rabbits.

The foraging behavior of wild rabbits (Oryctolagus cuniculus) is highly selective. They actively seek out energy-dense soft grass as their primary food source [59] and maintain necessary foraging activity to minimize the risk of predation [60]. Given the choice, rabbits prefer high-energy pellets over coarse forage [61, 62]. When consuming pellet feed, rabbits secrete more amylase, and the hardness of the food (through chewing) is the primary stimulus for saliva secretion [63]. Examples of dietary adaptation to high-starch diets during domestication are dogs and water buffalo, both of which had a mutation in the *AMY2B* [64, 65].

The coat color is an important domestication trait, and white rabbits are currently the most common domesticated rabbits. We compared breeds with white fur and breeds with colored fur, and discovered significant divergence in a partial region of the TYR gene. This gene is a crucial developmental locus that controls melanin production, and it has been extensively studied in animals such as humans [66], mice [67], and rabbits [68].

## Methods

### Sample selection and genome sequencing

In this study, a total of 167 rabbits were analyzed for genome resequencing from various regions. The samples included five local Chinese rabbit breeds, namely Laiwu Black rabbit (LWB; n = 20), Sichuan White rabbit (SCW; n = 20), Jiuyishan rabbit (JYS; n = 20), Fujian White rabbit (FJW; n = 18), and Minxishan Black rabbit (MXN; n = 14); three introduced breeds, namely Rex Rabbit (REX; n = 15), Angora Rabbit (AN; n = 20), and New Zealand White Rabbit (NZW; n = 10); one bred breed, namely Kangda Meat Rabbit (KD; n = 15); and representatives of two subspecies of wild rabbits (WILD; n = 15), namely Oryctolagus cuniculus cuniculus, primarily originating from the northern regions of Spain and southern parts of France, and Oryctolagus cuniculus algirus, primarily originating from the southern regions of Spain. The wild rabbits and New Zealand White rabbits were obtained from the NCBI database [69, 70], while the rest were collected for sequencing (Supplementary Table S1). Total genomic DNA was extracted and purified from rabbit ear tissues using the standard phenol/chloroform extraction method. For each sample, two paired-end libraries (500 bp) were constructed according to the manufacturer’s protocols (Illumina) and sequenced on the Illumina HiSeq 2500 sequencing platform. We sequenced each sample at a depth of 10X in order to reduce the false-negative rate of variants due to our strict filter criteria.

### Sequence alignment and variant calling

Raw sequencing reads were trimmed using fastp software [71] to filter out low-quality bases and sequences. The following filtering criteria were applied: removing reads with ≥10% unidentified nucleotides (N), >10 nucleotides aligned to the adapter, or of which >50% bases had Phred quality scores less than 5. High-quality reads were aligned against the rabbit reference genome (OryCun2.0) using bwa “BWA-MEM” algorithm [72]. Duplicate reads were removed from individual sample alignments using SAMtools (v.0.1.19) [73]. The genomic variants for each accession were then identified with the HaplotypeCaller module and the GVCF model using Genome Analysis Toolkit (GATK) software [74]. All the GVCF files were merged. We filtered variants both per population and per individual using the HaplotypeCaller module according to the stringent filtering criteria. The selection criteria are as follows: a) depth for individualC≥C4, b) genotype quality for individualC≥C5, c) minor allele frequency (MAF)C≥C0.05, d) with a missing rateC≤C0.1. The identified SNPs and InDels were further annotated with ANNOVAR [75], and divided into specific genomic regions and genes. To estimate the allele frequency of individual SNPs, we use the formula to calculate the absolute allele frequency difference (ΔAF) between each SNP in domestic and wild rabbits: ΔAF = abs (RefAFdom – mean (RefAFfrench + RefAFiberian)). Next, we categorize the SNPs by ΔAF in increments of 0.05 (i.e., ΔAF = 0-0.05, 0.05-0.10, etc., up to 0.95-1.00), and intersect these categorized SNPs with mammalian non-coding exons, UTRs, introns, and evolutionarily conserved non-coding elements [69, 76], the operation of transferring from hg18 to OryCun2 was conducted using the liftOver tool [77], for each ΔAF (allele frequency change) interval, a Python script was used to determine the proportion of SNPs that fall into non-coding exons, UTRs, introns, and conserved elements among 29 mammals through intersection.

### Population structure and phylogenetic analyses

The population structure was inferred using the program ADMIXTURE (v1.23) [78]. For each of the different subgroups (KC=C2–6), the population classification and the ancestry composition of each individual were simulated, and it was found that clusters maximized the marginal likelihood with K = 6 value. The phylogenetic tree analysis was performed using IQ-TREE software (v1.6.6) [79], based on the best model (GTRC+CFC+CASCC+CR7) selected by the Bayesian information criterion. The results were visualized by the the online tool EvolView (https://www.evolgenius.info//evolview/). PCA was performed using both GCTA software [80]. The population relatedness and migration events were inferred using TreeMix [81]. Nucleotide diversity (π) and fixation index (F_ST_) were calculated by vcftools [82] and pairwise genetic distance was calculated by Arlequin [83]. To study the genetic distance between different breeds, the squared correlation coefficient (r^2^) between pairwise SNPs was computed using the PopLDdecay [84] software. Parameters in the program were --MaxDist 500 --MAF 0.05 --Miss 0.1. The average r^2^ value was calculated for pairwise markers in a 500-kb window and averaged across the whole genome.

### Demographic history reconstruction

We used the Pairwise Sequentially Markovian Coalescent (PSMC) based on Single Nucleotide Polymorphism (SNP) distributions to infer the demographic history of both wild and domesticated rabbits [85]. According to previous research and suggestions, we adjusted the number of free atomic time intervals (-p option), the upper limit of time to the most recent common ancestor (-t option), and the initial value of r = θ/ρ (-r option) [86]. Moreover, we employed the SMC++ method, which can infer the effective population size history from hundreds of individuals and is more efficient than PSMC in recovering the history for very short time scales [87]. For PSMC/SMC++ analysis, scaling was performed using an average mutation rate (μ) of 1.25 × 10-8 per base per generation and a generation time (g) of 1 year.

### Mitochondrial haplotype analysis and gene flow analysis

The sequences were aligned using Mega 7.0 software [88], and DnaSP (6.12.03) was utilized to identify all haplotypes [89], polymorphic sites, and parsimony informative sites. We used Treemix (v1.13) conducted on the non-linkage SNP data [81], with the settings -root -bootstrap -k 1000 -m, and values for (−m) ranging from 1 to 30.

### Selective-sweep analysis

The F_ST_ and π were calculated using VCFtools with sliding windows of 50 kb, which had a 10 kb overlap between adjacent windows. The top 5% regions were identified as candidate selective regions, and genes within these regions were considered as candidates. All differentially expressed genes underwent GO functional annotation and KEGG pathway enrichment analysis using clusterProfiler [90]. The term or pathway was considered significantly enriched when the false discovery rate (FDR) was less than or equal to 0.05.

### Data availability

The sequencing data used for analysis is available at NCBI (PRJNA1011829)

